# BLADE-R: streamlined RNA extraction for molecular diagnostics and high-throughput applications

**DOI:** 10.1101/2025.03.27.645479

**Authors:** Anam Tajammal, Samuel Haddox, Shafaque Zahra, Robert Cornelison, Adelaide Ohui Fierti, Hui Li

## Abstract

Efficient nucleic acid extraction and purification are crucial for cellular and molecular biology research, yet they pose challenges for large-scale clinical RNA sequencing and PCR assays. Here, we present BLADE-R, a magnetic bead-based protocol that simplifies the process by combining cellular lysis and nucleic acid binding into a single step, followed by a unique on-bead rinse for nuclease-free separation of genomic DNA and RNA. The Agilent TapeStation and RT-qPCR analyses show that RNA extracted from HEK293T cell line using BLADE-R outperforms the TRIzol protocol in terms of time and cost. RNA sequencing reveals no differences in sequence quality or gene count variance between samples processed with BLADE-R and those processed with TRIzol followed by RNA kit clean-up. Additionally, BLADE-R outperformed TRIzol in RNA extraction from frozen tissue and whole blood samples, as confirmed by RT-qPCR. Our protocol can be adapted to a 96-well plate format, enabling RNA purification of up to 96 human blood samples in less time than a single-sample traditional extraction. Using BLADE-R in this format, we confirmed minimal well-to-well contamination in RNA purification, cDNA synthesis, and PCR. Therefore, our novel BLADE-R protocol, suitable for both low and high-throughput formats, is effective even in limited-resource settings for preparing clinical samples for PCR and sequencing assays. Thus, our new BLADE-R technique works well even in low-resource environments to prepare clinical samples for PCR and sequencing experiments. It can be adapted for both low- and high-throughput formats.

## Introduction

Friedrich Miescher first precipitated DNA from the nuclei of leukocytes in 1862 using acetic or hydrochloric acid, giving rise to the term “nucleic acid” (NA) [1]. As most molecular biology assays begin with the separation of DNA, RNA, and/or protein there has been a worldwide, concerted effort to increase the quality and yield of biomolecular extractions for almost two centuries. However, as Polymerase Chain Reaction (PCR) and DNA/RNA sequencing-based assays become more clinically relevant, there is increasing pressure for more streamlined, and high-throughput, lab methods for NA extraction and processing to meet clinical demands. Selecting the right NA extraction technique is crucial since it affects the precision of the molecular diagnosis [2]. Currently, phenol-based liquid-phase and solid-phase silica column techniques are widely used for nucleic acid extraction due to their ease and cost-effectiveness. Despite varying in approach, all methods share the same goals: cell lysis, nuclease inactivation, and efficient biomolecule isolation (3).

Liquid phase separation methods typically utilize an acidic phenol, guanidine isothiocyanate, and chloroform-based protocol to provide high-quality extraction [4]. Phenol is used to denature and precipitate proteins while dissolving hydrophobic cellular components into the organic phase. When mixed with chloroform, the density of the phenolic organic phase is increased, preventing a mixture with the aqueous phase containing the hydrophilic NA [5]. The aqueous phase must be carefully removed and mixed with isopropanol or ethanol to precipitate NAs into an often-unobservable pellet. Air-drying of the pellet is used to remove contamination and the pellet is reconstituted with Tris-EDTA (TE) buffer or nuclease-free water [3]. In contrast to liquid phase, solid phase separation methods offer a distinct approach to nucleic acid purification by using a solid phase matrix to bind nucleic acids. Materials such as agarose [6], diatomaceous earth [7], glass particles [8], or silica [9] silica can serve as the matrix.

As a popular alternative to commercially available RNA extraction reagents, such as TRIzol^®^ (Thermofisher, Waltham, MA), solid phase separation kits that implement a silica membrane in a microcentrifuge tube sized column, such as Qiagen’s RNeasy kit (ThermoFisher, Waltham, MA), are commonly used these days. While suitable for high-throughput processing with extensive lab equipment, column-based kits are not cost-effective in low-resource settings [10], [11]. Magnetic bead-based nucleic acid purification is popular for its efficiency, ease of use, and suitability for high-throughput workflows [12]. Studies have shown magnetic beads effectively extract nucleic acids from various biological materials, including whole blood [13], serum [14], sputum [15], tissue, and urine [16]. While the magnetic bead-based methods effectively address scalability concerns, the integration of sample pretreatment and a distinct lysis step is cumbersome in high throughput settings particularly in the absence of automation [17], [18].

DNA purification is straightforward as DNA-targeting nucleases are easily denatured with phenol alone [3], and RNA presence doesn’t affect most downstream analyses. On the contrary, extraction of pure, high-quality RNA for downstream analysis is of particular concern as RNases happen to be more resilient to inactivation [5], [19], and downstream gene expression analysis protocols will be affected by DNA contamination [20]. Currently, there are no commercially accessible high-throughput magnetic bead-based RNA extraction kits that produce DNA-free RNA without requiring a nuclease treatment. Therefore, the primary focus of this study is RNA isolation approaches with a focus on high-throughput clinical PCR assays that use a minimal amount of starting material. This study introduces an innovative protocol termed BLADE-R **(beads lysis and DNA elimination for RNA extraction)** consisting of a single step for cell lysis and nucleic acid binding, followed by “on-bead” wash steps and a nuclease-free gDNA removal rinse to achieve a high-throughput, high efficiency, and rapid extraction of RNA even in low resource settings.

## Results

### Establishment of the lysis bead reagent and protocol

To optimize one-step lysis beads for the BLADE-R method, we tested several lysis buffer combinations with magnetic beads, including mixes of SDS, DTT, Tris(2-carboxyethyl) phosphine (TCEP), GTC, Triton X-100, and Tween 20 in TE buffer. The combination of DTT, 2M GTC, Tween 20, and (PEG-8000) in TE buffer provided the best lysis results and optimal RNA concentration and purity. Therefore, we established this protocol using that combination for lysis beads and Wash buffer 1.

Combining a well-established bead-based NA purification method with lysis reagents and on-bead DNA removal enabled the isolation of RNA directly from cells as shown in Figure 1A. DTT and GTC was premixed with the magnetic beads and binding buffer and stored as a single reagent, termed lysis beads. Wash buffer 1, which has the same formulation as the lysis bead reagent but without any magnetic beads, serves to continue lysis and resuspend the beads after the initial lysis. Wash buffer 2, which has the same formulation as Wash buffer 1 but without DTT and GTC, serves to remove lysis components along with any remaining cellular debris. Other than the DNA removal method, the remaining protocol, consisting of ethanol rinses and NA elution. We streamlined the extraction process to efficiently remove DNA from RNA without the need for additional purification steps.

**Figure 1:** Comparison of RNA extraction protocols using BLADE-R (beads lysis and DNA elimination for RNA extraction) and the traditional TRIzol method. (**A**) Workflow of RNA extraction using BLADE-R, detailing each step of the protocol and the corresponding time required. (**B**) Workflow of RNA extraction using the TRIzol method, detailing each step and the corresponding time required. Figure created with BioRender.com.

RNA obtained from this BLADE-R protocol can be directly employed in downstream applications, such as RT-qPCR/TaqMan assays and RNA sequencing. Conversely, RNA extracted from TRIzol requires either DNase treatment or a DNA-removing column clean-up [21]. Even in a low-throughput context, the BLADE-R protocol provides significant time reductions (>300%) when compared to a standard protocol such as TRIzol as shown in Figure 1B. The cost savings, however, are more dramatic as reagents in BLADE-R are less than one-third the price of TRIzol per reaction further detailed in Supplementary Figure 1 and Supplementary Table 1.

### LiCl rinse for on-bead removal of DNA

Utilizing LiCl, a substance reported to selectively precipitate RNA over DNA (ThermoFisher, Waltham, MA) [22], we were able to prepare a bead rinse that removes DNA from the beads while leaving RNA precipitated. Figure 2A illustrates the workflow and highlights the efficacy of using LiCl for DNA removal during RNA extraction. Unlike other methods that require a separate DNA removal step with DNase or additional bead cleanup after RNA purification, our approach provides RNA that is immediately ready for downstream applications. To evaluate the efficacy of the DNA-removing LiCl on-bead rinse, we employed qPCR to quantitate relative intron *GAPDH* normalized by exon *GAPDH* as an indicator of DNA contamination and assessed cDNA prepared from BLADE-R extractions with no DNA removal compared to RNA from BLADE-R protocols using LiCl DNA removal, DNases treatment, and both in combination. Figure 2B indicates that 5M LiCl DNA removal performs equal to DNase, and samples lacking LiCl, or DNase treatment displayed the highest genomic DNA contamination. This underscores the efficacy of LiCl in removing genomic DNA. Although LiCl is traditionally used for RNA precipitation, we have demonstrated a novel approach by utilizing LiCl in the form of an on-bead rinse to remove DNA before the final RNA elution. Unlike traditional methods that require RNA cleanup after elution, this technique streamlines the RNA extraction process, making it more rapid and yielding ready-to-use RNA for any downstream applications without the need for further cleanup. To create conditions that would increase DNA solubility in the LiCl rinse, we evaluated room temperature (RT) and 4°C LiCl rinses with higher vs. lower concentrations of LiCl. Optimization of the DNA-removing LiCl rinse revealed that room temperature 0.2M LiCl rinse may be used as substitute of the cold 5M LiCl (Supplementary Figure 2).

**Figure 2:** LiCl effectively removes DNA during RNA extraction. (**A**) Workflow illustrating DNA removal steps in various RNA extraction methods. (**B**) Assessment of DNA contamination in U87 cells via qPCR analysis, comparing LiCl rinse, DNase treatment, a combination of both, and no treatment. Figure 2A was created with BioRender.com.

### RNA quality assessment and Gene Expression Analysis in cell lines

RNA was purified from 1.2x10^6^ HEK293T cells using TRIzol, and BLADE-R. RNA yield and purity was assessed through a nanodrop spectrophotometer and RNA integrity was assessed using Agilent TapeStation. Although samples extracted from TRIzol had higher RNA concentration, BLADE-R samples showed higher 260/280 and 260/230 ratios (Figure 3A) suggesting purer RNA is extracted using the BLADE-R. All samples extracted from the BLADE-R showed RNA integrity number of 10 (Figure 3B). For assessment of RNA quality for downstream applications, RT-qPCR was performed to amplify a highly expressed housekeeping gene *GAPDH,* and a lowly expressed gene *AVIL* in HEK293T cell lines. The comparative analysis of cycle threshold (Ct) values, as depicted in Figure 3C, indicates lower Ct values of *GAPDH* in samples extracted from BLADE-R in contrast to those extracted from TRIzol. Amplification plots and melt curves for *GAPDH* underscore the uniformity in performance between RNA extracted from BLADE-R and TRIzol (Figures 3D and 3E).

**Figure 3:** RNA quality assessment and Gene Expression Analysis in cell lines. (**A**) Nanodrop analysis showing RNA concentration and purity from HEK293T cells, with B1-3 representing samples extracted using BLADE-R and T1-3 representing samples extracted using TRIzol. (**B**) RNA integrity assessment via Agilent TapeStation, demonstrating superior RNA integrity in samples extracted using BLADE-R. (**C**) Mean Ct values showing *GAPDH* expression in HEK293T cells from RNA extracted using BLADE-R and TRIzol, analyzed by SYBR Green qPCR assay. (**D**) Amplification plots for *GAPDH* expression in HEK293T cells. (**E**) Melt curve plots for *GAPDH* (**F**) Mean Ct values showing *AVIL* expression in HEK293T cells from RNA extracted by BLADE-R and TRIzol. (**G**) Amplification plots for *AVIL* gene in HEK293T cells. (**H**) Melt curve plots for *AVIL* expression in HEK293T cells.

Furthermore, gene expression analysis for *AVIL* gene showed lower Ct values in samples extracted from BLADE-R as compared to TRIzol (Figure 3F). Amplification plots and melt curve plots depict the amplification efficiency of *AVIL* gene between both methods (Figure 3G and 3H).

These findings collectively support the notion that RNA extracted from BLADE-R is equivalent to that obtained from TRIzol, with the former exhibiting a favorable characteristic of lower Ct values, signifying superior amplification efficiency.

### RNA-sequencing analysis of purified RNA

RNA sequencing is a powerful downstream application of RNA extracted from various biological sample types. In addition to quantifying gene expression, RNA-Seq data enables the discovery of new transcripts, identification of alternatively spliced genes, and detection of allele-specific expression [23]. To demonstrate the efficacy of the BLADE-R extraction method for RNA-seq, we extracted RNA from human Astrocytes (HA) cell line using BLADE-R and TRIzol methods. Paired-end 150 bp Illumina HiSeq was performed comparing the samples purified with TRIzol followed by Qiagen RNeasy kit clean-up to BLADE-R method. FastQC is used to identify problems in sequencing files before alignment and/or after trimming. FastQC output showed no differences in sequence quality between extraction methods as shown in Figure 4A. Alignment was carried out using the traditional aligner, STAR [24], and the pseudoaligner, Kallisto [25], [26]. After counts were tabulated, we assessed the overall differences between replicate samples and a sample prepared with our BLADE-R method. Pearson correlation plots of sample comparison showed similar differences between all samples. RNA samples extracted from BLADE-R and TRIzol show identical correlation scores of 0.96 (Figure 4B). Looking at transformed count totals and overall expression densities, there were no significant differences between samples prepared with BLADE-R or with the traditional TRIzol method as shown in Figure 4C. These results suggest that RNA purified with BLADE-R can be used to produce high-quality RNA sequencing library preps without introducing any detectable sequencing bias when compared to RNA purified with TRIzol followed by Qiagen kit RNA cleanup.

**Figure 4:** RNA sequencing analysis of RNA extracted using BLADE-R and TRIzol. (**A**) FastQC output showing comparable mean quality scores of sequences and demonstrating similar per-sequence quality scores for RNA extracted by BLADE-R and TRIzol (followed by Qiagen kit RNA cleanup) methods. (**B**) Correlation matrix indicating that RNA sequencing results from samples extracted using BLADE-R are highly correlated with those extracted using TRIzol (followed by Qiagen kit RNA cleanup) method. (**C**) Distribution of transformed data and density plot showing similar data distribution for RNA samples extracted using both methods.

### Evaluation of RT-qPCR amplification efficiency of purified RNA from human frozen tissue

We also assessed the quantity and quality of RNA extracted from human frozen liver samples, focusing on two key genes, *GAPDH* and *Actin*. We utilized different RNA isolation methods: TRIzol alone, TRIzol followed by magnetic beads clean-up, and BLADE-R. We found that RNA extracted using TRIzol was the most concentrated among all the methods, whereas RNA isolated with BLADE-R yielded RNA with the least concentration (Figure 5A). Furthermore, we investigated the influence of RNA purity on gene expression analysis by comparing the cycle threshold (Ct) values of *GAPDH* and *Actin*. Lower Ct values indicate higher expression levels and better RNA quality. Our results showed that Ct values decreased progressively from TRIzol to TRIzol followed by bead purification, and finally to the BLADE-R. This trend indicates that RNA extracted with the BLADE-R had the highest RNA quality, resulting in more accurate and reliable gene expression data (Figures 5B and 5C). The BLADE-R method emerged as particularly advantageous, yielding RNA with the highest purity and consequently enhancing the reliability of our gene expression results.

**Figure 5:** RT-qPCR analysis of RNA extracted from frozen human tissue samples using BLADE-R, TRIzol, and RNA clean-up (magnetic beads) after TRIzol extraction. (**A**) Assessment of RNA quantity and purity extracted from frozen human liver tissues using the three methods. (**B**) Mean Ct values for *GAPDH* amplification from RNA extracted using three different approaches. (**C**) Mean Ct values for *Actin* amplification from RNA extracted using three different approaches.

### Evaluation of RT-qPCR amplification efficiency of purified RNA from human blood

Human blood has long been a vital diagnostic resource due to its continuous interaction with all body tissues. Biobanks routinely collect blood specimens for medical applications, scientific research, and diagnostics [27], [28]. Due to its wide application in diagnostics and gene expression profiling, small volumes of blood are also useful for RNA extraction. For downstream studies, we aimed to establish RNA extraction protocol from lower blood volume. We purified RNA using BLADE-R and TRIzol using just 20 µl of blood samples. We compared Ct values of *GAPDH* and *SRRM2* genes amplified from RNA extracted from whole blood using both methods. Ct values of *GAPDH* were lower in samples extracted from beads than samples extracted from TRIzol (Figure 6A). We also amplified *SRRM2* gene, where we observed statistically significant differences in Ct values. BLADE-R showed lower Ct values (high amplification efficiency) than the TRIzol (Figure 6B). Amplification plots and melt curve plots show amplification efficiency and specificity respectively (Figures 6C and 6D). Furthermore, we extracted RNA from blood clots collected in golden top tubes, used for serum testing containing clots at the bottom (these samples were stored at 4°C for 28 days). RT-qPCR amplification of *GAPDH* and *SRRM2* genes demonstrated that samples extracted using BLADE-R were still suitable for gene expression analysis (Supplementary Figure 3).

**Figure 6:** RT-qPCR analysis of RNA extracted from human clinical blood samples using BLADE-R and TRIzol method. (**A**) Mean Ct values showing GAPDH expression in RNA extracted from human blood samples using BLADE-R and TRIzol. (**B**) Mean Ct values showing *SRRM2* expression in RNA extracted from human blood samples using BLADE-R and TRIzol. Unpaired student t-test was performed as statistical test using, p-value = 0.0031. (**C**) Amplification plots for *GAPDH* and *SRRM2*. (**D**) Melt curve plots for *GAPDH* and *SRRM2*.

### Establishment of High-Throughput Protocol

For processing of large-scale clinical patient cohorts, we modified BLADE-R protocol to a high-throughput 96-well plate format. We prepared 96-well plates containing wash buffers, LiCl rinse, and ethanol rinse in bulk and stored them for a week. After mixing sample and lysis beads reagent in a 96-well plate, a 96-tipped magnet was utilized to move magnetic beads from lysis plate to previously prepared wash and rinse plates (Figure 7A). This eased the RNA extraction from 96 samples in almost as little time as needed for a single sample extraction from other traditional methods. We initially tested the high-throughput protocol using cultured K562 cells distributed across a 96-well plate, with nuclease-free water as no-template controls (Figure 7B). Maintaining the same plate format through reverse transcription, melt curve analysis revealed that only K562 sample wells exhibited the correct Tm, indicating no detectable cross-contamination between the sample and water wells.

**Figure 7:** High-throughput BLADE-R extraction from K562 cells and clinical blood samples. (A) Workflow of high-throughput RNA extraction from cell culture or blood samples. (B) Mean Ct values showing *SRRM2* gene expression from RNA extracted from K562 cells and water (control) in a 96-well format. (C) Mean Ct values showing *GAPDH* expression from RNA extracted from clinical blood samples in a 96-well format. Figure 7A was created with BioRender.com.

Once we confirmed minimal well-to-well contamination, we repeated the protocol using just 13 µl of clinical blood for each sample. As shown in Figure 7C, most of the clinical blood samples had a *GAPDH* Ct value of less than 20 when using 14 µl of RNA input from the lysis bead elution for cDNA synthesis and 2 µl of cDNA input per well for qPCR reaction. These results suggest that the 96-well format protocol provides a rapid and cost-effective solution for RT-qPCR clinical blood assays and requires less than 20 µl of blood to perform multiple assays, eliminating the need for additional blood draws.

## Discussion

Combining lysis reagents and on-beads genomic DNA removal before the final RNA elution with an already well-established magnetic bead protocol, RNA extraction can be performed rapidly on cell culture, blood samples, and homogenized tissue samples [13], [17]. Furthermore, storage of the lysis and binding components along with the magnetic beads as a single reagent has proven to be stable for periods exceeding 6 months in 4LJ (data not shown). In this study, we have adapted this approach to high-throughput RNA isolation for RT-PCR and RNA-seq-based clinical assays. Moreover, this protocol holds great potential to enhance the cost-effectiveness of RNA isolation across both research and clinical laboratory settings.

BLADE-R protocol provides a more rapid alternative to TRIzol that yields RNA of equal or better quality. Other similar bead protocols (*Quick*-RNA Miniprep Kit, Zymo Research, MagMAX™ *mir*Vana™ Total RNA Isolation Kit, ThermoFisher) utilize DNase and therefore require additional cleanup, which in most cases is another round of the bead binding, wash, and elution steps. Our BLADE-R protocol employs LiCl for DNA removal, reducing bead reagent usage by half, and nearly reducing the time of the protocol by half as another round of bead clean-up is not necessary. We have demonstrated that the BLADE-R can effectively extract high-quality RNA from cell culture samples with ease, even without repeating the wash and rinse steps shown in Fig 1A. However, processing blood samples could be more challenging in some cases.

A key advantage of the BLADE-R protocol is its high throughput capacity. While well-to-well contamination is a common concern in 96-well plate, careful execution of the BLADE-R protocol minimizes this issue. We have shown that expensive automation-based approaches are not necessary for quick and efficient RNA extractions, and a manual technique that utilizes a 96-well pipette tip magnet and 96-well plates pre-aliquoted with reagent, wash, and rinse could achieve extraction and sample preparation times comparable to automation-based approaches. Our protocol in 96-well plate format can be adapted for automation by utilizing a 96-well plate magnet like already established bead-based automated approaches with the advantage of the on-bead nuclease-free gDNA removal that reduces time and reagent usage. The reagent and protocol described here are the first to describe a nuclease-free gDNA removal on-bead rinse for magnetic bead-based RNA purification that is ideal for evaluating RNAs in clinical samples such as blood or frozen tissue samples. We believe that the 96-well plate format of the described BLADE-R protocol will grant clinics access to molecular biology assays to diagnose a large range of diseases and infections without a major buy-in cost or specialized core facilities.

This study presents a novel, cost-effective, and high-throughput RNA extraction using lysis reagents and magnetic beads, which surpasses traditional techniques in both efficiency and RNA quality. The protocol is adaptable for various sample types and laboratory settings, providing significant benefits for research and clinical applications. By enabling rapid and high-quality RNA isolation, this method holds great promise for advancing molecular diagnostics and therapeutic research.

## Materials and Methods

In this study, we used the following recipes for the BLADE-R method consisting of lysis beads reagent, wash buffers, 5M LiCl, and 70% Ethanol solutions for RNA extraction from various biological samples including clinically discarded blood and tissues as provided in Table 1.

**Table 1.** Recipe of the buffers used for lysis bead method for RNA extraction.

|  |  |
| --- | --- |
| Lysis Beads reagent | 7.5mM dithiothreitol (DTT), 2 M Guanidine Thiocyanate (GTC), 6 g PEG-8000, 1M NaCl, 500 µl of 1 M Tris-HCl pH 8.0, 100 µl of 0.5 M EDTA, Tween 20 27.5 µl, 0.5 ml magnetic beads dissolved in 50 ml of nuclease-free water. |
| Wash Buffer 1 | 7.5mM DTT, 2 M GTC, 6 g PEG-8000, 1M NaCl, 500 µl of 1 M Tris-HCl pH 8.0, 100 µl of 0.5 M EDTA, and 27.5 µl Tween 20 dissolved in 50 ml of nuclease-free water. |
| Wash Buffer 2 | 6 g PEG-8000, 1M NaCl, 500 µl of 1 M Tris-HCl pH 8.0, 100 µl of 0.5 M EDTA, and 27.5 µl Tween 20 dissolved in 50 ml of nuclease-free water. |

### Ethics Statement

The use of the human clinical blood samples and frozen tissues was approved by the University of Virginia’s Institutional Biosafety Committee (IBC) and Biosafety Office under protocol HSR# 13310. All experimental protocols were approved by University of Virginia’s Institutional Biosafety Committee (IBC) and Biosafety Office. Relevant regulations and institutional policies were strictly followed.

### Chemical reagents

All reagents used in this study are of analytical grade and did not require any further purification. Guanidine Thiocyanate (cat#J65104.30), DTT (Lot#01336167), NaCl (cat#194848), and LiCl (cat#449045000) were purchased from Thermo Scientific, USA. Sera-Mag™ Carboxylate-Modified Magnetic Beads (Lot#17264624) were purchased from Cytiva, USA. EDTA (0.5 M) (Lot#2085657) was purchased from Invitrogen, USA. PEG-8000 (Lot#113H0309) from molecular sigma biology and Tris-HCl from Fisher Scientific were already available in the lab.

### RNA extraction from cell culture using BLADE-R and TRIzol

Cell pellets were resuspended in 100 µl of sterile PBS, and 600 µl of lysis beads were added. After vortexing, samples were left for 10 minutes. Samples were immobilized on a magnetic rack, and supernatant was removed. Tubes were taken off the magnetic rack, and 600 µl of Wash buffer 1 was added to each, followed by vortexing. After 1 minute, samples were vortexed and immobilized on magnetic rack. The supernatant was discarded, wash with buffer 1 was performed. Next, Wash buffer 2 was used to perform two bead washes. After discarding the supernatant of second wash, sample tubes were kept on magnetic rack for subsequent LiCl and ethanol rinse steps. 5M LiCl solution was added to cover the magnetic beads and left for a minute before removing the LiCl (twice). After rinsing with LiCl, the tube was filled with 70% ethanol and allowed to sit for 2 minutes. Ethanol was removed, and this step was repeated once more. After removing ethanol for the second time, tubes were kept on magnetic rack to air dry the magnetic beads. Once dried, the sample tubes were removed from magnetic rack, and 20 µl of DNase & RNase free water was added to resuspend the magnetic beads and concentration was measured.

To extract RNA using TRIzol, cell pellet was resuspended in 1 ml of TRIzol solution. Samples were vortexed and left at room temperature for 10 minutes and 0.2 ml chloroform per 1 ml of TRIzol reagent was added, followed by vigorous shaking for 15 seconds and incubation for 2-3 minutes. After centrifugation at maximum speed at 4°C for 15 minutes, the top layer was collected and transferred to a clean tube, 1μl of RNase-free glycogen and 0.5 ml of RNase-free isopropanol was added. After incubation at -80°C for 30 minutes, the samples were centrifuged at maximum speed for 10 minutes at 4°C. The supernatant was removed, and the RNA pellet was washed twice with 1 ml of 75% ethanol and centrifuged at maximum speed for 5 minutes. The RNA pellet was air-dried for approximately 10 minutes. The RNA was resuspended in 20 μl of RNase-free water and incubated for 10 minutes at 55°C to aid in dissolution. To extract RNA from blood using TRIzol 20 µl of buffy coat was resuspended in 180 µl of 1X PBS and 500 µl TRIzol was added, and the above-stated method for TRIzol was exactly repeated.

RNA extraction from liver was performed using three different methods. The first one was using TRIzol reagent. After the initial step of crushing 100 mg liver tissue with liquid nitrogen using a mortar and pestle into a fine pulverized powder, TRIzol was used for sample lysis followed by the above stated method. The second method uses RNA extraction from TRIzol followed by beads purification. 1.8X of SeraPure beads were added to purified RNA followed by 5M LiCl treatment (twice). The beads-RNA complexes were captured using a magnetic stand and contaminants were removed by subsequent washing with 80% EtOH followed by elution. Lastly, RNA was extracted using the BLADE-R. After crushing the tissue using liquid nitrogen, 600 µl of lysis beads were added. After homogenization, the lysate was processed, and RNA was extracted using BLADE-R protocol.

### RNA quality and integrity assessment

Concentration in ng/µl, 260/230, and 260/230 ratios were measured for each sample by using NanoDrop™ 2000 Spectrophotometer. RNA integrity was assessed using Agilent bioanalyzer/tape station by Genome Analysis and Technology Core, RRID:SCR_018883 of the University of Virginia.

### Agarose gel electrophoresis

1% Agarose gel was prepared by dissolving 0.5 g of agarose in a 50 ml TAE buffer and 5 μl EtBr was added. 5 µl of Meridian Bioscience HyperLadder 100bp was loaded to assess the size. Bluejuice 10x loading buffer by Invitrogen, USA was added to each sample. Gel was run for 30 minutes at 90 volts. Gel was visualized by the gel documentation system after exposure to UV light.

### RT-qPCR

cDNA Synthesis was performed using Thermo Scientific Verso cDNA synthesis kit with oligo(dT) and 2 µg of RNA as per manufacturer’s instructions. Quantitative PCR was performed using 2X Universal SYBR Green from ABclonal cat#RK21203 as per the manufacturer’s instructions. Amplification was performed for 40 cycles, each at 95°C for 15 s and 60°C for 1 min, using the Quantstudio 3 Real-Time PCR system (Thermo Fisher Scientific). For high-throughput clinical blood RNA, ABscript Neo RT Master Mix for qPCR with gDNA remover kit from Abclonal cat#RK20433 was used as per manufacturer’s instructions. The list of primers used is presented in Supplementary Table 2. All measurements were performed in triplicates.

### Experimental RNA sequencing workflow

RNA was extracted from U87 cells using the TRIzol method as per manufacturer’s instruction and the BLADE-R method as described above. Samples extracted from TRIzol samples were subjected to column clean-up before sending for RNA sequencing while samples extracted from BLADE-R were not subjected to any clean-up method. Samples were sent to Novogene for paired-end 150 bp sequencing with samples averaging ∼25 million reads.

The preliminary analysis started with assessing the overall read quality using FastQC (http://www.bioinformatics.babraham.ac.uk/projects/fastqc), compiled with MultiQC [29]. After passing QC, reads were aligned to the human genome (Ensembl GRCh38.v110) using Kallisto [25], [26] with the -GC bias flag, and bootstrapping set to 100 for downstream use of Sleuth [26]. After alignment with Kallisto abundances were passed to DESeq2 [30], [31] using tximport [32] and differential expression analysis was performed. To test for batch effects induced by the two extraction methods we variance stabilized and used the removebatcheffect function from limma [33] and Combat-seq from the SVA package [34]. To check for batch effects from unknown sources we ran Combat-seq with default settings. Since Kallisto uses a pseudoalignment approach, we also did a second pipeline swapping Kallisto for STAR [24]. We performed an extended quality control analysis on the raw reads using iSEE [35], and fastp [36], [37] to check for any unanticipated issues. Hierarchical clustering, k-means clustering, and other advanced visualizations were performed in R, using: PCAexplorer [38], [39], and IDEP [40]. Fastp output reports were used to easily summarize reads by Phred score and compare between extraction techniques.

### High-throughput RNA extraction of clinical blood samples

10 million K562 cells were resuspended in 2 ml PBS. 20 µl of cell suspension was added to each well containing a sample and RNAse-free water was added for control wells. 120 µl of lysis beads were added to all wells and a multichannel pipette was used to resuspend the samples thoroughly. The plate was kept at rest for 15 minutes. Using Emerther Esmart magnetic pipette, beads were transferred to a plate containing 180 µl of Wash buffer 1. The beads were resuspended in Wash buffer 1 and kept for rest for 5 minutes. The beads were moved to another plate with Wash buffer 2 and the process was repeated as for first washing. Magnetic beads were transferred to the plate containing LiCl, samples were allowed to soak in LiCl without removing the pipette. After 2-3 minutes beads were moved to a 70% ethanol plate for soaking. After soaking in ethanol, beads were dried and resuspended in 20 µl of RNAse-free water, followed by cDNA synthesis and RT-qPCR.

For high-throughput RNA extraction from blood, we utilized a rack of clinical blood samples that had been stored at 4°C for 72 hours. Due to extended shelf time, natural separation occurred within the tubes, resulting in distinct layers of plasma, white blood cells, and red blood cells. Plasma layer was carefully avoided, and the middle layer of the buffy coat was used as a sample. We collected 13 µl of buffy coat from each blood sample and resuspended in 7 µl of 1X PBS in each well of the 96-well plate. 180 µl of lysis beads were added in each well and by using multichannel pipette samples were subjected to lysis. After lysis, the method stated above for high-throughput RNA extraction from cell culture was used, with each step being done twice.

### Data Analysis

GraphPad prism10 was used to analyze and run statistical tests on qPCR data in this study.

## Data availability

The supporting data for this study, have been deposited in the gene expression omnibus repository (GEO) under accession number GSE278095.

## Acknowledgments

We thank Dr. Lindsay Bazydlo for providing clinical blood samples and the Biorepository and Tissue Research Facility (BTRF) at the University of Virginia for providing clinical tissue samples.

## Funding

HL is supported by NIGMS R01GM132128, NCI R01CA245905, R01CA240601, and R01CA269594.

## Author contributions

H.L., A.T., & S.H., conceived the idea, interpreted the data, and wrote the manuscript; A.T., performed experiments; R.C., performed the bioinformatic analysis; S.Z., conducted experiments on frozen tissues; A.F., contributed to the manuscript preparation. All authors read the final version of the manuscript and agreed to its publication.

## Conflict of interests

The authors declare that they have no conflict of interest.

## Notes

### Competing Interest Statement

The authors have declared no competing interest.

### Summary of Updates

1. Table 1: Recipe of the buffers used in the protocol corrected The final concentration of NaCl has been corrected from 5 M to 1 M. 2. In the results section, the cell line used for the RNA-sequencing analysis of purified RNA has been corrected from U87 to Human Astrocytes (HA).

