## Supplementary figures and images for "BLADE-R: streamlined RNA extraction for molecular diagnostics and high-throughput applications" (from another version of this preprint: version 1, 2025-03-29)

### supplementary 1

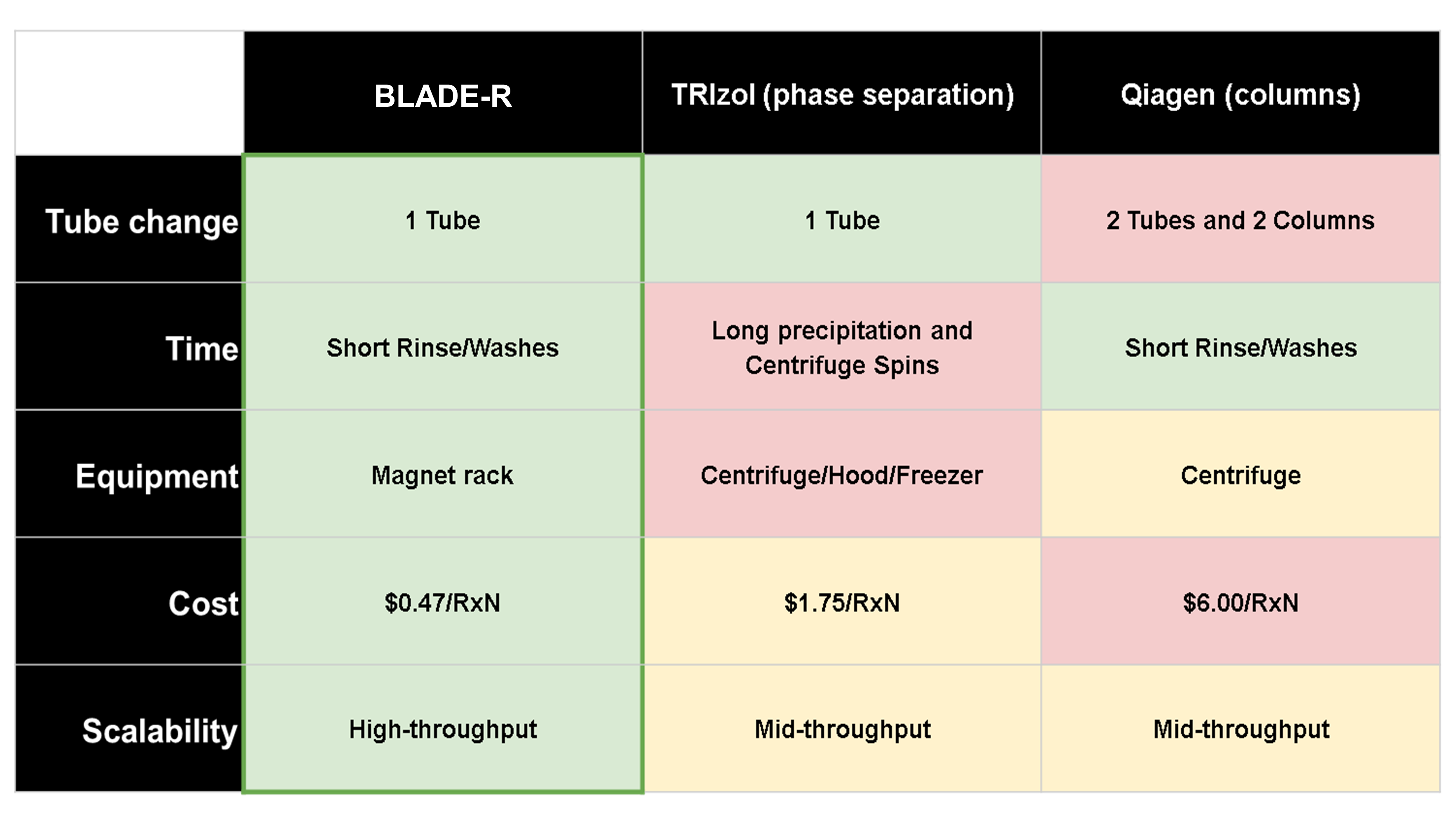

### supplementary 2

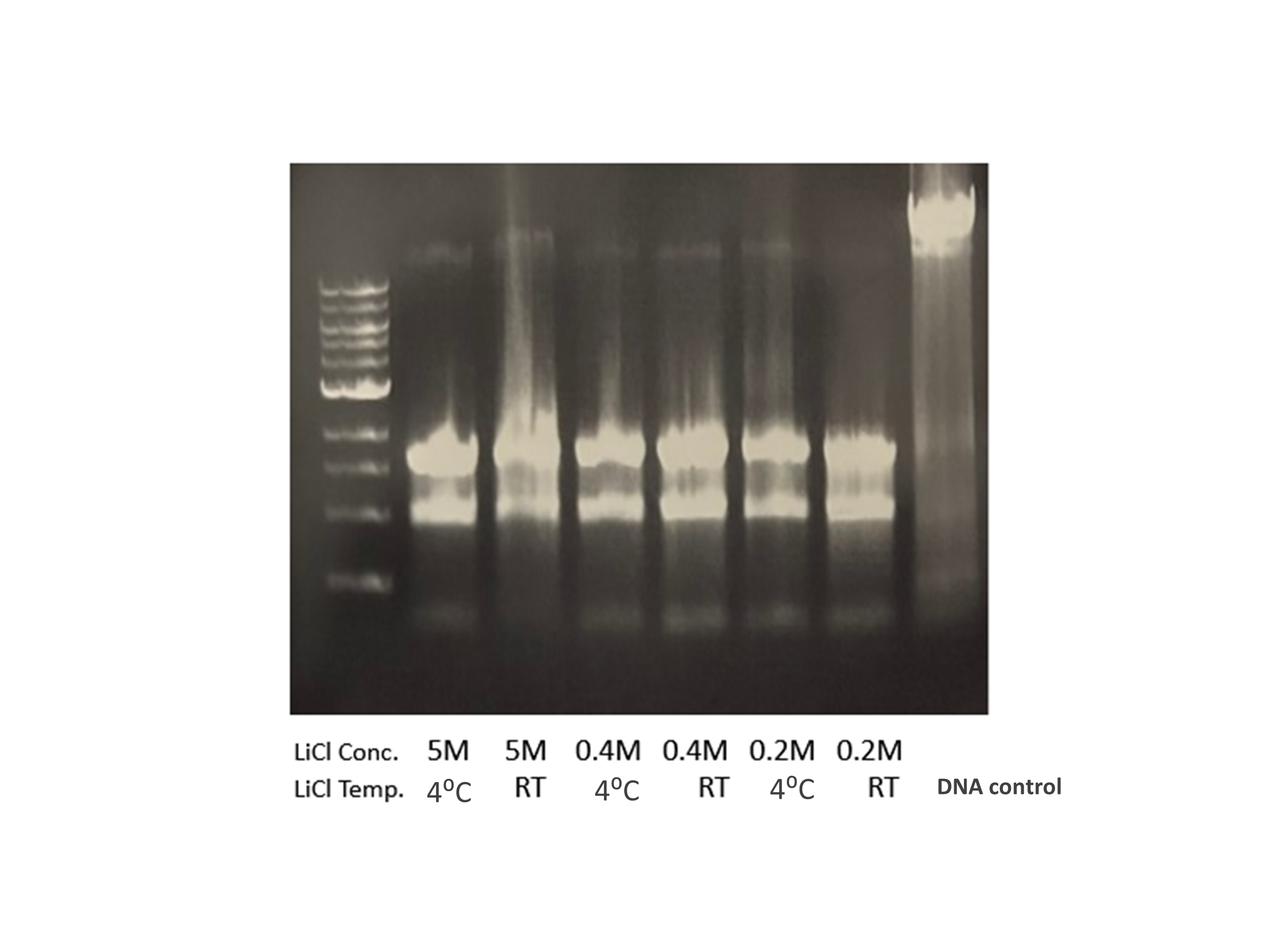

### supplementary fig 3

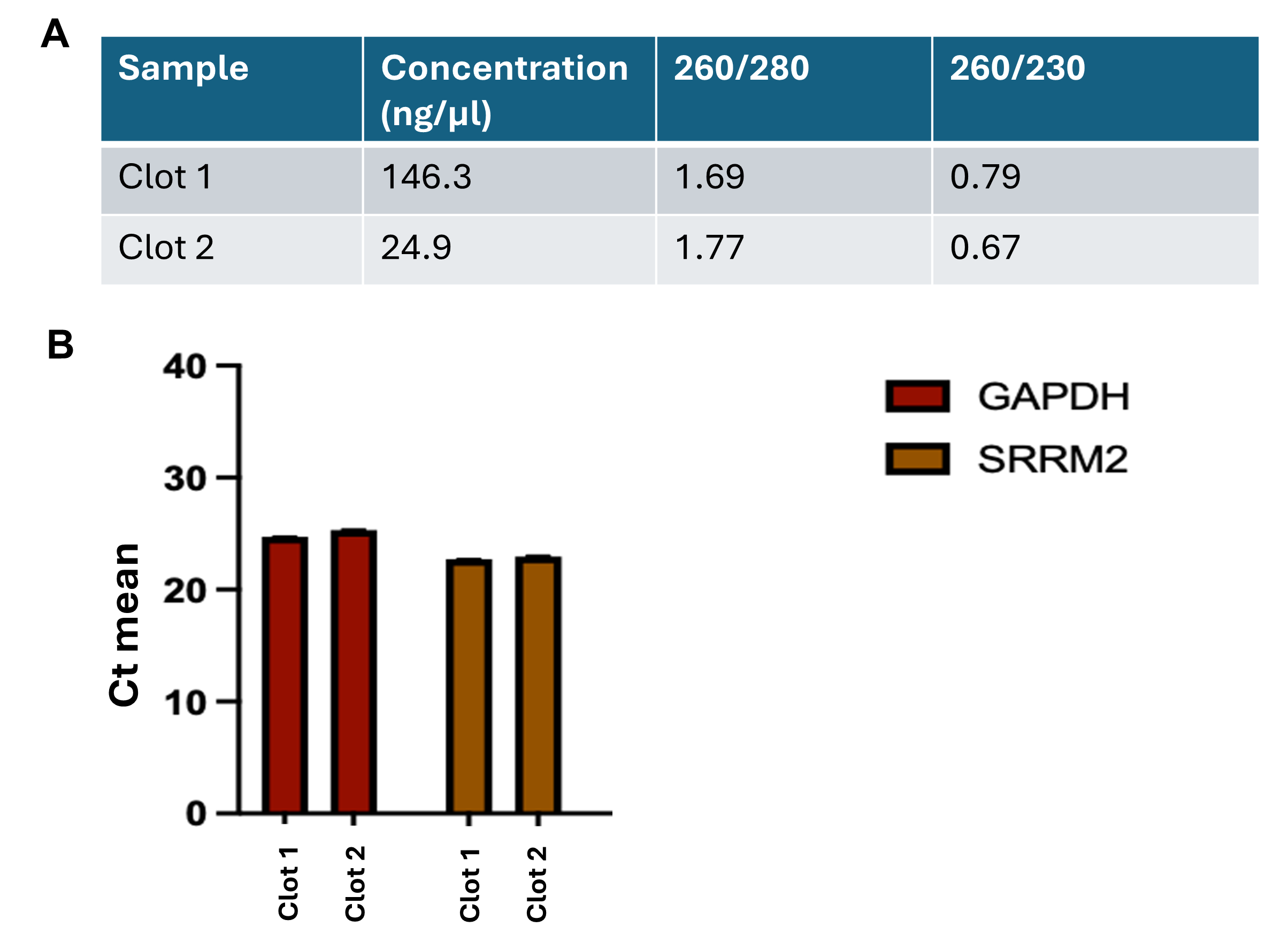
