## supplementary Tables for "BLADE-R: streamlined RNA extraction for molecular diagnostics and high-throughput applications" (from another version of this preprint: version 1, 2025-03-29)

Supplementary Table 1: Comparison of buffer reagents and per reaction cost of BLADE-R and TRIzol.

| **Reagents** | **Cost ($)** | **Quantity** | **Unit(s)** | **Amount per 50ml** | **Lysis buffer** | **Wash buffer1** | **Wash buffer2** | **LiCl** |
| --- | --- | --- | --- | --- | --- | --- | --- | --- |
| Cytiva speedbead magnetic carboxylate | 500 | 15 | ml | 0.5 | 16.66666667 | NA | NA | NA |
| PEG-8000 | 174 | 1000 | g | 6 | 1.044 | 1.044 | 1.044 | NA |
| 0.5 M EDTA pH 8.0 | 59.5 | 500 | ml | 0.1 | 0.0119 | 0.0119 | 0.0119 | NA |
| 1.0 M Tris pH 8.0 | 23 | 100 | ml | 0.5 | 0.115 | 0.115 | 0.115 | NA |
| Tween 20 | 203 | 4000 | ml | 0.0275 | 0.001395625 | 0.001395625 | 0.001395625 | NA |
| 5 M NaCl | 513 | 10000 | ml | 10 | 0.513 | 0.513 | 0.513 | NA |
| LiCl | 282 | 500 | g | 4.2 | NA | NA | NA | 2.3688 |
| GITC | 234.5 | 500 | g | 11.86 | 5.56234 | 5.56234 | NA | NA |
| DTT | 213.5 | 5 | g | 0.057 | 2.4339 | 2.4339 | NA | NA |
|  |  |  |  | cost per 50ml | 26.34820229 | 9.681535625 | 1.685295625 | 2.3688 |
| **Method** | **Cost ($)** | **Quantity** | **Unit** | **Amount of each reagent per sample** | **Cost per sample ($)** |  |  |  |
| TRIzol | 439 | 200 | ml | 0.6 | **1.32** |  |  |  |
| BLADE-R | 40.083 | 50 | ml | 0.6 | **0.47** |  |  |  |

**Supplementary Table 2:** Primer details used in the study.

| Primers | Sequence | Figures |
| --- | --- | --- |
| Intron GAPDH F | CTTACCTGTGCTCCCACTCC | Figure 2 |
| Intron GAPDH R | ACCCCTTCTCTAAGTCCCTCC | Figure 2 |
| AVIL F | TGCGAACTCAGGCAGAGCACTA | Figure 3 |
| AVIL R | CAGTGTTCTCGCTGCCATCACA | Figure 3 |
| Actin F | CACCATTGGCAATGAGCGGTTC | Figure 5 |
| Actin R | AGGTCTTTGCGGATGTCCACGT | Figure 5 |
| SRRM2 F | GGCAGCTCTTTTGATCCTCAGC | Figures 7 |
| SRRM2 R | GTGAGCGAGAACTGCTAGACTC | Figures 7 |
| Exon GAPDH F | ACCCACTCCTCCACCTTTGA | Figures 2,3,5,7 |
| Exon GAPDH R | CTGCTGCTGTAGCCAATTCGT | Figures 2,3,5,7 |
